# Male accessory gland factors increased sex pheromone titres in the glands of *Spodoptera litura* female moths by inhibiting calling behavior

**DOI:** 10.1101/872150

**Authors:** Jin Xu, Min-Rui Shi, Da-Ying Fu, Hong Yu, Peng Chen, Hui Ye

**Affiliations:** Yunnan Academy of Biodiversity, Southwest Forestry University, Kunming 650224, China; Yunnan Academy of Forestry, Kunming 650201, China; School of Life Sciences, Yunnan University, Kunming 650091, China

**Author notes:** Correspondence: Peng Chen,; Hui Ye.

**Keywords:** Sex pheromone, Mating, MAG secretions, Calling behavior, Sex pheromone release, Neural control

## Abstract

Moths are the most widely studied example of pheromones in animals. However, little is known about the mechanism of intrasexual and mating-related intersexual regulation of pheromone production and release in female moths. Our previous studies in *Spodoptera litura* found that mating induced a higher sex pheromone titre in the pheromone gland (PG) and mating or male accessory gland (MAG) extract suppressed female calling behavior. We therefore hypothesize that the inhibition of female calling behavior by mating or MAG factors likely suppresses the release of sex pheromones and thus results in a higher pheromone titre in the PG. To test this hypothesis, in the present study, we introduced an artificial calling behavior suppression treatment by gently knocking on and shaking the testing boxes contained moths once every 10 minutes. Results show that this treatment significantly increased pheromone titres in virgin or saline injected virgin females, and the increase rates are similar to those of mating and MAG extract treated ones. These results have suggested that the increase of sex pheromone titer in the female PG after mating in *S. litura* is due to the inhibition of female calling behavior by MAG factors. Moreover, results of this study also suggest that female calling behavior is positively correlated to pheromone release and likewise, the calling behavior and sex pheromone release in *S. litura* females are directly under the neural control, and modulated by molecular and environmental factors.

## INTRODUCTION

To date, the sex pheromones for more than 2000 insects species have been identified, including more than 600 moth species (Allison & Cardé, 2016, The Pherobase, 2019). Accordingly, great progress has been achieved in recent years on the internal mechanisms of sex pheromone biosynthesis and release in insects (Jurenka, 2017). Based on these findings, sex pheromone-based mass trapping and mating disruption have been successfully used in forecasting and controlling many pest insects (Srinivasan et al., 2015, Allison & Cardé, 2016).

In moths, sex pheromones usually are produced in a pheromone gland (PG) associated with the ovipositor in females, and pheromone components are emitted during a specific calling behavior (Percy-Cunningham & MacDonald, 1987). Previous studies have indicated that calling behavior and pheromone release are modulated by neural, hormonal or a combination of both (reviewed in Solari et al., 2007), but these mechanisms are not unequivocally demonstrated in Lepidopterans.

Previous studies in more than twenty insect species (most are moths) generally found that mating can induce the termination of sex pheromone production and a lower sex pheromone titre in the PG (reviewed in Lu et al., 2017). Kingan et al. (1995) identified a male accessory gland (MAG) produced pheromonostatic peptide (PSP) in *Helicoverpa zea*, which produced a short-term (few hours) effect in depleting females of pheromones. Other studies also demonstrated that the MAG-produced sex-peptide (SP) and its receptor (SPR) in females, which were found in *Drosophila* and function in the regulation of postmating behavior (Chen et al., 1988, Yapici et al., 2008), may also exist in moths and can inhibit female calling behavior and pheromone synthesis as well (Nagalakshimi et al., 2007, Li et al., 2014). Moreover, some studies also suggested that the presence of sperm in the spermatheca may trigger the release of a signal, via the ventral nerve cord (VNC), to suppress pheromone production in female moths (Giebultowicz et al., 1991, Delisle et al., 2000). Molecular studies revealed that sex pheromone production in female moths is regulated by the Pheromone Biosynthesis Activating Neuropeptide (PBAN) (Raina & Kempe, 1990). Mating seems will not affect the the mRNA expression of *PBAN* (Fodor et al., 2017), while may inhibit the release of PBAN to the hemolymph and will result in a low pheromone titre in the PG (Nagalakshimi et al., 2007). Nevertheless, the possible links between MAG factors and PBAN are still unclear.

Interestingly, our recent study (Lu et al., 2017) has found an opposite result in *Spodoptera litura* where mating resulted in a higher sex pheromone titre in the PG. Our other previous studies in *S. litura* also showed that sex pheromone production in females is also regulated by PBAN (Lu et al., 2015), and mating, injection of MAG extract and RNA interference of *SPR* significantly suppressed female calling behavior and receptive to males (Li et al., 2014, Yu et al., 2014). Therefore, we hypothesize that the suppression of female calling behavior by mating or MAG factors likely inhibits the release of sex pheromones and thus will result in a higher sex pheromone titre in the PG (Lu et al., 2017).

Above studies have implied that the mechanism of intrasexual and mating-related intersexual regulation of pheromone production and release still a challenging but fascinating mystery. In the present study, therefore, we set up and conducted a series of treatments and experiments in *S. litura* to test above developed two hypotheses: (1) mating (with sperm and MAG factors), as well as MAG factors (without sperm), will induce an increase of sex pheromone titre in the PG, and (2) mating induced sex pheromone titre increase in the PG is due to the suppression of calling behavior (which inhibits pheromone release) by MAG factors.

The common cutworm moth, *Spodoptera litura* (Lepidoptera: Noctuidae) is a key agricultural pest worldwide. Four sex pheromone components have been identified for *S. litura* (Tamaki Y, 1973, Tamaki et al., 1976): Z9,E11-14:Ac (A), Z9,E12-14:Ac (B), Z9-14:Ac (C) and E11-14:Ac (D), with a strict 100:27:20:27 (A:B:C:D) ratio in PG.

## MATERIALS AND METHODS

### Insects

*Spodoptera litura* larvae were reared on an artificial diet (Li et al., 1998), and adult moths were fed a 10% honey solution under a 14:10 h light:dark photoperiod, at 26°C and 60–80% relative humidity. Only 1-d-old virgin moths with an average body weight (within Mean±1SD) (Lu et al., 2017) were used in this study.

### Effect of MAG extract, mating and calling behavior suppressing on sex pheromone titre in the PG

Eight female treatments were set up: (1) virgin females were injected with 6μl MAG extract (equivalent to one male MAG tissue) (MG), (2) virgin females were injected with 6μl MAG extract, and then their calling behavior were artificially suppressed by gently knocking on and shaking the testing boxes contained moths once every 10min (our previous observation found that *S. litura* females are quite sensitive and this treatment can effectively prevent female calling) (MG+S), (3) virgin females were injected with 6μl saline (Sl), and (4) virgin females were injected with 6μl saline, and then their calling behavior were artificially suppressed as above (Sl+S), (5) virgin females (Vr), (6) virgin females with their calling behavior were artificially suppressed as above (Vr+S), (7) mated females (Mt), and (8) mated females with their calling behavior were artificially suppressed as above (Mt+S). MAG extract preparation and injections followed the methods outlined in Yu et al. (2014). Injections were conducted and matings were allowed during 4-5h of the second scotophase after emergence. After injections and matings, females were kept individually in plastic boxes (18cm long, 10cm wide, 6cm high) and the artificial calling behavior suppressing were conducted or not according to the treatments as mentioned above until the time of sampling. The PGs were sampled (four PGs were used as a replicate) at 16 and 24h after treatment (injections or matings) and the sex pheromone titres (n = 6) were measured by GC using the same protocol outlined in (Lu et al., 2017).

### Statistical analysis

Data were analyzed using an ANOVA followed by Fisher’s LSD test. All analyses were conducted using SPSS 16.0. The rejection level was set at *α* < 0.05. All values reported are Means±SE.

## RESULTS

ANOVA revealed that sex pheromone titres in the PGs were significantly different between treatments both at 16h (*F*_7,40_=4.99, *P*<0.0001 for compound A; *F*_7,40_=4.94, *P*<0.0001 for compound B; *F*_7,40_=3.96, *P*<0.001 for compound C; and *F*_7,40_=9.14, *P*<0.0001 for compound D) and 24h (*F*_7,40_=6.56, *P*<0.0001 for compound A; *F*_7,40_=2.45, *P*<0.05 for compound B, *F*_7,40_=4.72, *P*<0.001 for compound C; and *F*_7,40_=3.42, *P*<0.001 for compound D) after treatment (Fig. 1).

**Fig. 1.**
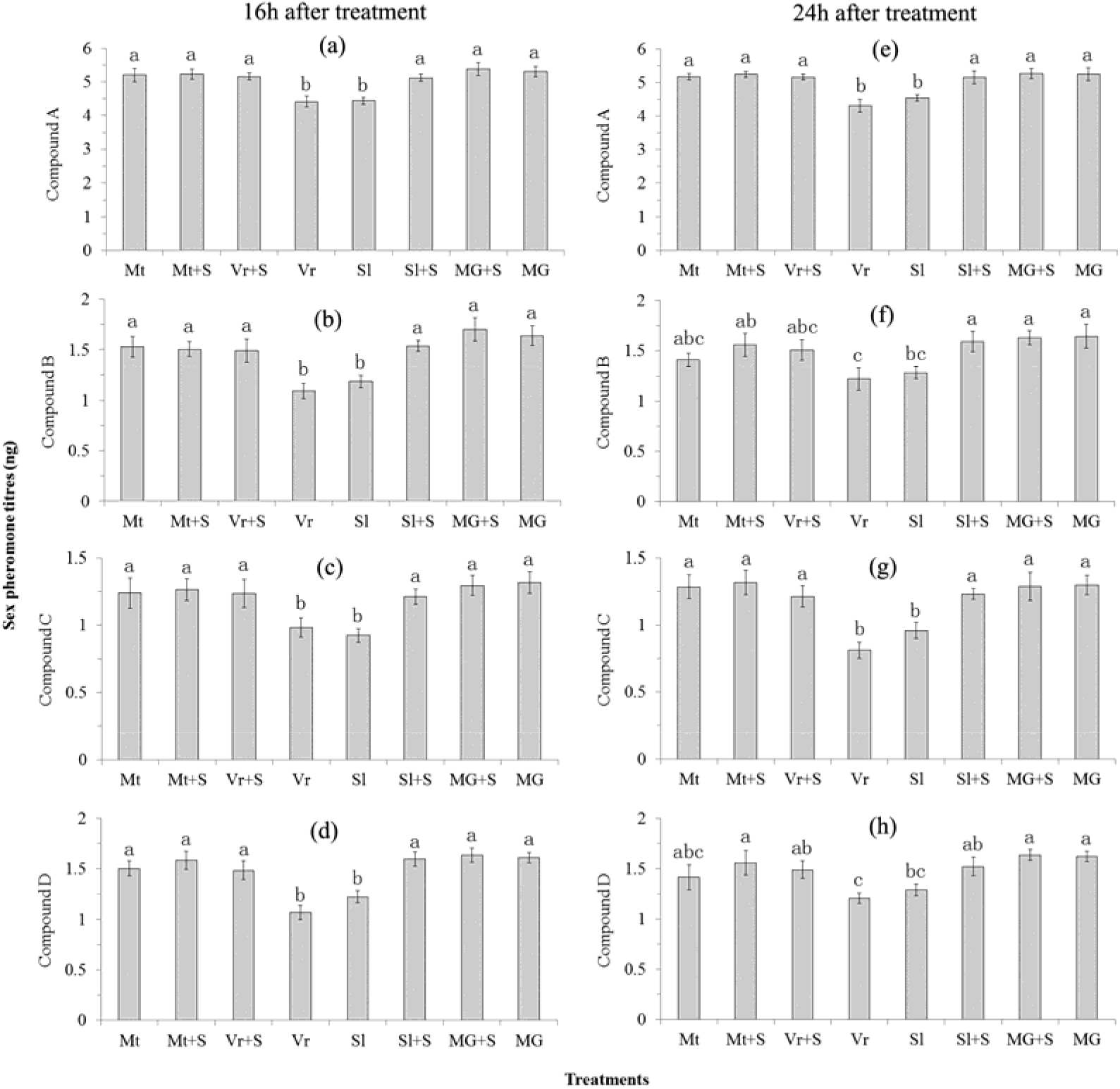
Effect of MAG extract, mating and calling behavior suppressing on sex pheromone titres in the PGs of *S. litura* females. (a) to (d) were measured at 16h after treatment while (e) to (h) were measured at 24h after treatment. For each compound of 16 or 24h after treatment, the bars with different letters are significantly different (*P* < 0.05).

Post hoc LSD test showed that females of treatment MG, MG+S, Sl+S, Vr+S, Mt and Mt+S have significantly higher sex pheromone titres than that of treatment Vr and Sl (*P* < 0.05) in most cases (Fig. 1 a-e, g) except for compound B and D at 24h after treatment (Fig. 1 f & h).

## DISCUSSION

In our previous study (Lu et al., 2017), we compared sex pheromone titres in the PG between mated and virgin *S. litura* females at 4, 8, 16 and 24h after mating and found mating significantly increased pheromone titres at 16 and 24h after mating. In the present study, we further found that either mating or MAG extract (without sperm) significantly increased sex pheromone titres in the PG of *S. litura* females in most cases at 16 and 24h after mating or MAG extract injection (Fig. 1 a-e, g). It is still unknown whether sperm also has any effect on sex pheromone production and release (Giebultowicz et al., 1991, Delisle et al., 2000) in *S. litura*, but the result of the present study (Fig. 1) has suggested that MAG factors should be the main trigger for the increase of sex pheromone titre in mated females.

Based on the evidence that MAG extract injection or RNA interference of the SPR gene can suppress female calling behaviors and matings in *S. litura* (Li et al., 2014, Yu et al., 2014), we hypothesize that the suppression of female calling behavior by MAG factors is likely to shut off the release of sex pheromone and thus will induce a higher sex pheromone titre in the PG. To test this hypothesis, we introduced an artificial calling behavior suppression treatment try to inhibit sex pheromone release in *S. litura* females. Results show that this treatment significantly promoted pheromone titres in virgin or saline injected virgin females, and the increase rates are similar to that of mating and MAG extract treated females (Fig. 1). These results suggest that the mating induced increase of sex pheromone titre in the PG of female *S. litura* is due to the suppression of sex pheromone release by MAG factors. Moreover, there is no significant difference was found between treatment Mt and Mt+S, as well as between MG and MG+S, suggesting that the artificial pheromone release suppression treatment had no stress on these females in terms of sex pheromone titres.

Calling behavior are believed to under the control of the ventral nerve cord (VNC) (Tang et al., 1987) and the caudalmost terminal abdominal ganglion (TAG) (Itagaki & Conner, 1987), and hormonal (such as juvenile hormone) and environmental (such as light) factors (Delisle & McNeil, 1987, Liang & Schal, 1994). During calling, the abdomen of female *S. litura* protrudes between the wings with the tip everted, and the PG exposed and expanded (visible at this time) (Li et al., 2012). In *S. litura*, female calling behavior only occur at night, not during the daytime (Li et al., 2012). Our previous observation also found that *S. litura* females are quite sensitive and very light vibration will interrupt their calling behavior, i.e. the abdomen returned to the normal post and the PG retracted (invisible at this time) (J. Xu, unpubl. data); the behavior of ceasing calling and pheromone release when facing risks may have evolved in *S. litura* to evade the attack of natural enemies as its sex pheromone may also attract parasitoids and predators as well (e.g. Powell et al., 1993). In the present study, we successfully suppressed female calling behavior (no calling behavior was observed) and sex pheromone release in treatment Sl+S and Vr+S (Fig. 1) artificially by gently knocking on and shaking the boxes contained the testing moths once every 10 minutes, suggesting that the calling behavior and sex pheromone release also are under the neural control.

However, the molecules and mechanisms between mating and the suppression of pheromone release still are unclear. Based on the results of the present study and other studies on PBAN, PSP, SP and SPR (reviewed in Lu et al., 2017), we propose that the suppression of female calling behavior and sex pheromone release after mating may be due to the effect of the SP-like peptide in MAG fluids, which warrants further study.

## Acknowledgement

Research reported here was supported by projects from the Joint Special Project of Yunnan Province for Basic Agricultural Research (2018FG001-002), and the National Natural Science Foundation Program of P.R. China (31760635; 31560606; 31660208).

## REFERENCES

Allison, J.D. and Cardé, R.T. 2016. Pheromone communication in moths: evolution, behavior and application, University of Caifornia Press, Oakland, California.

Chen, P.S., Stummzollinger, E., Aigaki, T., Balmer, J., Bienz, M., et al. 1988. A male accessory gland peptide that regulates reproductive behavior of female *Drosophila melanogaster*. Cell, 54, 291–298.

Delisle, J. and Mcneil, J.N. 1987. Calling behaviour and pheromone titre of the true armyworm *Pseudaletia unipuncta* (Haw.) (Lepidoptera: Noctuidae) under different temperature and photoperiodic conditions. J. Insect. Physiol., 33, 315–324.

Delisle, J., Picimbon, J. and Simard, J. 2000. Regulation of pheromone inhibition in mated females of *Choristoneura fumiferana* and *C. rosaceana*. J. Insect. Physiol., 46, 913–921.

Fodor, J., Koblos, G., Kakai, A., Karpati, Z., Molnar, B.P., et al. 2017. Molecular cloning, mRNA expression and biological activity of the pheromone biosynthesis activating neuropeptide (PBAN) from the European corn borer, *Ostrinia nubilalis*. Insect Mol. Biol., 26, 616–632.

Giebultowicz, J.M., Raina, A.K., Uebel, E.C. and Ridgway, R.L. 1991. Two-step regulation of sex-pheromone decline in mated gypsy moth females. Arch. Insect. Biochem., 16, 95–105.

Itagaki, H. and Conner, W.E. 1987. Neural control of rhythmic pheromone gland exposure in *Utetheisa ornatrix* (Lepidoptera: Arctiidae). J. Insect. Physiol., 33, 177–181.

Jurenka, R. 2017. Regulation of pheromone biosynthesis in moths. Curr. Opin. Insect Sci., 24, 29–35.

Kingan, T.G., Bodnar, W.M., Raina, A.K., Shabanowitz, J. and Hunt, D.F. 1995. The loss of female sex pheromone after mating in the corn earworm moth *Helicoverpa zea*: Identification of a male pheromonostatic peptide. P. Natl. Acad. Sci. USA., 92, 5082–5086.

Li, C., Yu, J.-F., Lu, Q., Xu, J., Liu, J.-H., et al. 2014. Molecular characterization and functional analysis of a putative sex-peptide receptor in the tobacco cutworm *Spodoptera litura* (Fabricius, 1775) (Lepidoptera: Noctuidae). Aust. Entomol., 53, 424–431.

Li, C., Yu, J.-F., Xu, J., Liu, J.-H. and Ye, H. 2012. Reproductive rhythms of the tobacco cutworm, *Spodoptera litura* (Lepidoptera: Noctuidae). GSTF J. BioSci., 2, 25–29.

Li, G., Chen, Q. and Pang, Y. 1998. Studies of artificial diets for the beet armyworm, *Spodoptera exigua*. Acta Sci. Nat. Univ. Sunyatseni, 4, 1–5.

Liang, D. and Schal, C. 1994. Neural and hormonal regulation of calling behavior in *Blattella germanica* females. J. Insect. Physiol., 40, 251–258.

Lu, Q., Huang, L.-Y., Liu, F.-T., Wang, Xia-Fei, Chen, P., Xu, J., et al. 2017. Sex pheromone titre in the glands of *Spodoptera litura* females: circadian rhythm and the effects of age and mating. Physiol. Entomol., 42, 156–162.

Lu, Q., Huang, L.Y., Chen, P., Yu, J.F., Xu, J., et al. 2015. Identification and RNA interference of the Pheromone Biosynthesis Activating Neuropeptide (PBAN) in the common cutworm moth *Spodoptera litura*. J. Econ. Entomol., 108, 1344–1353.

Nagalakshimi, V.K., Applebaum, S.W., Azrielli, A. and Rafaeli, A. 2007. Female sex pheromone suppression and the fate of sex-peptide-like peptides in mated moths of *Helicoverpa armigera*. Arch. Insect. Biochem., 64, 142–155.

Percy-Cunningham, J.E. and Macdonald, J.A. 1987. Biology and ultrastructure of sex pheromone-producing glands, in: G.D. Prestwich & G.J. Blomquist (Eds.) Pheromone biochemistry. Academic Press, Inc, Orlando, Fl.

Powell, W., Hardie, J., Hick, A.J., Holler, C., Mann, J., et al. 1993. Responses of the parasitoid *Praon volucre* (Hymenoptera, Braconidae) to aphid sex-pheromone lures in cereal fields in autumn - implications for parasitoid manipulation. Eur. J. Entomol., 90, 435–438.

Raina, A.K. and Kempe, T.G. 1990. A pentapeptide of the C-terminal sequence of PBAN with pheromonotropic activity. Insect. Biochem., 20, 849–851.

Solari, P., Crnjar, R., Spiga, S., Sollai, G., Loy, F., et al. 2007. Release mechanism of sex pheromone in the female gypsy moth *Lymantria dispar*: a morpho-functional approach. J. Comp. Physiol. A, 193, 775–785.

Srinivasan, R., Lin, M.-Y., Su, F.-C., Yule, S., Khumsuwan, C., et al. 2015. Use of insect pheromones in vegetable pest management: successes and struggles, pp. 231–237. in: A.K. Chakravarthy (Ed.) New Horizons in Insect Science: Towards Sustainable Pest Management. Springer, New Delhi.

Tamaki Y, N.H., Yushima T 1973. Sex pheromone of *Spodoptera litura* (F.) (Lepidoptera: Noctuidae): isolation, identification and synthesis. Appl. Entomol. Zool., 8, 200–203.

Tamaki, Y., Osawa, T., Yushima, T. and Noguchi, H. 1976. Sex pheromone and related compounds secreted by the virgin female of *Spodoptera litura* (F.) Jpn. J. Appl. Entomol. Z, 20, 81–86.

Tang, J.D., Charlton, R.E., Cardé, R.T. and Yin, C.-M. 1987. Effect of allatectomy and ventral nerve cord transection on calling, pheromone emission and pheromone production in *Lymantria dispar*. J. Insect. Physiol., 33, 469–476.

The Pherobase 2019. The Pherobase: Database of insect pheromones and semiochemicals [Internet].

Yapici, N., Kim, Y.J., Ribeiro, C. and Dickson, B.J. 2008. A receptor that mediates the post-mating switch in *Drosophila* reproductive behaviour. Nature, 451, 33–37.

Yu, J.F., Li, C., Xu, J., Liu, J.H. and Ye, H. 2014. Male accessory gland secretions modulate female post-mating behavior in the moth *Spodoptera litura*. J. Insect. Behav., 27, 105–116.

